# Wax “tails” enable planthopper nymphs to self-right midair and land on their feet

**DOI:** 10.1101/2024.04.15.589523

**Authors:** Christina L. McDonald, Gerwin T. Alcalde, Thomas C. Jones, Ruby Ana P. Laude, Sheryl A. Yap, M. Saad Bhamla

**Affiliations:** George W. Woodruff School of Mechanical Engineering, Georgia Institute of Technology, Atlanta, GA 30332, USA; School of Chemical and Biomolecular Engineering, Georgia Institute of Technology, Atlanta, Georgia, United States; Institute of Weed Science, Entomology, and Plant Pathology, College of Agriculture and Food Science, University of the Philippines Los Baños, College, Laguna, 4031, Philippines; Department of Entomology, College of Agriculture, University of Southern Mindanaos, Kabacan Cotabato, Philippines

**Keywords:** wax, planthopper nymph, aerial righting, self-righting, aerial dynamics

## Abstract

The striking appearance of wax ‘tails’ — posterior wax projections on planthopper nymphs — has captivated entomologists and naturalists alike. Despite their intriguing presence, the functional roles of these structures remain largely unexplored. This study leverages high-speed imaging to uncover the biomechanical implications of these wax formations in the aerial dynamics of planthopper nymphs (*Ricania sp*.). We quantitatively demonstrate that removing wax tails significantly increases body rotations during jumps. Specifically, nymphs without wax projections undergo continuous rotations, averaging 4.3 *±* 1.9 per jump, in contrast to wax-intact nymphs, who narrowly complete a full rotation, averaging only 0.7 *±* 0.2 per jump. This suggests that wax structures effectively counteract rotation through aerodynamic drag forces. These stark differences in body rotation correlate with landing success: nymphs with wax intact achieve a near perfect landing rate of 98.5%, while those without wax manage only a 35.5% success rate. Jump trajectory analysis reveals transitions from parabolic to Tartaglia shapes at higher take-off velocities for wax-intact nymphs, illustrating how wax structures assist nymphs in achieving stable, controlled descents. Our findings confirm the aerodynamic self-righting functionality of wax tails in stabilizing planthopper landings, advancing our understanding of the complex interplay between wax morphology and aerial maneuverability, with broader implications for the evolution of flight in wingless insects and bioinspired robotics.

## Introduction

### Diversity of wax production in insects

Insect wax, a variable mixture of true waxes and other organic substances, often forms a body coating or develops into ornate structures when produced in substantial amounts [1]. Notably, sap-feeding insects such as scales, woolly aphids, whiteflies, psyllids, and planthoppers produce conspicuous wax structures [1, 2]. Among them, planthoppers (Fulgoromorpha) stand out as “the most remarkable wax producers among insects” [3]. Planthoppers in tropical environments can produce wax strands reaching up to 75 cm [4]. While female adults typically produce wax to cover their eggs, nymphs of several species also contribute to the diversity of wax formations, illustrating the prevalence of wax across different life stages (Figure 1) [5], [6]. Despite this diversity, the mechanisms behind the formation of these intricate wax structures remain largely unexplored.

**Fig. 1.**
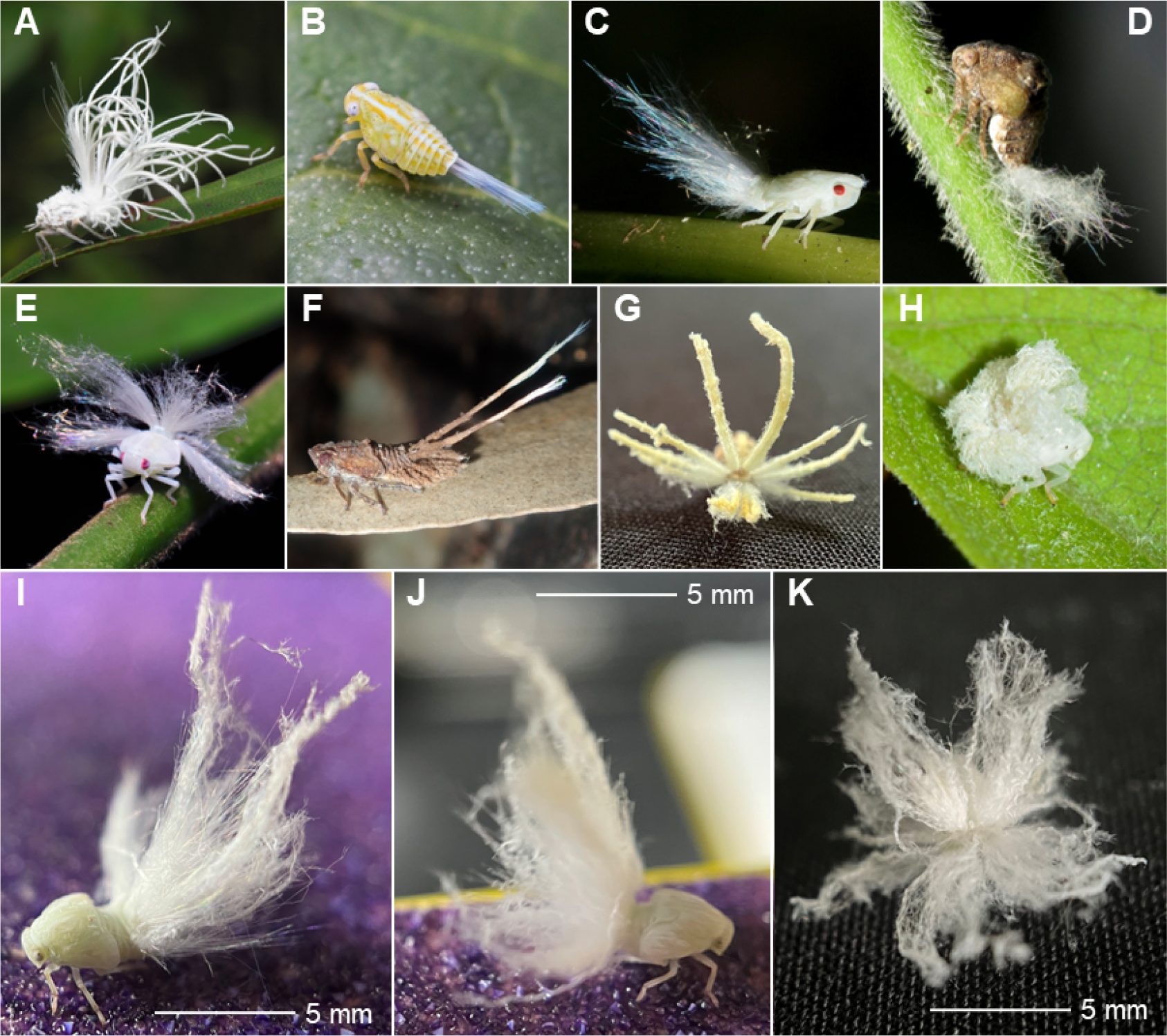
Planthopper nymphs produce a diversity of wax structures: A. A Flatid nymph from Cambodia [14]; B. An Issid nymph from the United States [15];C. A Flatid nymph from Peru [16]; D. An Acanaloniid nymph from the United States [17]; E. A Ricaniid nymph from Singapore [18]; F. An Eurybrachid nymph from Australia [19]; G. A Ricaniid nymph from the Philippines; H. A Ricaniid nymph from Taiwan [20]; I-K. *Ricania sp*. nymphs used in this study. Images A–F, H are unmodified and used under a Creative Commons Attribution License (CC BY or CC BY-NC).

Wax pores, mainly found on the abdomen, function as molds for shaping secreted wax. The diversity in wax pore morphology across species contributes to the variety of wax structures that planthopper nymphs produce [7]. The common appearance of wax ‘tails’, where wax accumulates at the nymph’s posterior, highlights this diversity [8]. The widespread occurrence and common posterior localization of these structures prompts a deeper investigation into their roles: What functions do wax tails serve?

### Multifaceted functions of wax

Insect wax performs several speculated and confirmed functions, providing benefits like microclimate control through desiccation resistance, UV radiation protection, and flood defense (due to the hydrophobicity of wax) [9, 3, 10]. Additionally, wax can deter predators through crypsis (camouflage) or act as a physical barrier against predators and parasites [7, 3, 11, 10, 2]. It may also protect against contamination from honeydew, the sticky, sugary excrement of planthoppers that is prone to fungal attacks [8].These multifaceted roles highlight the importance of wax in insect

survival and adaptability; however, while these functions are well-documented in various insect taxa, specific evidence for planthoppers remains limited [12]. The differentiation between wax coatings on the body surface and more morphologically complex wax formations, like the wax tails, seen in planthopper nymphs, raises questions about their functions. Considering the link between animal morphology and locomotion strategy [13], investigating the relationship between wax and the biomechancial performance of planthopper nymphs, especially their jumping capabilities, may offer new insights into the adaptive significance of these structures.

### Jumping and aerial righting

Planthoppers rank among the fastest jumpers in the insect world [21], suggesting a significant aerial phase post-take-off for both winged adults and wingless nymphs. For wingless animals, jumping and falling often leads to chagnes in body orientation. To counteract this, many animals employ active and passive strategies for mid-air reorientation, known as aerial righting [22]. The benefits aerial righting range from landing in positions conducive to further jumps to achieving targeted landing [23, 24, 25]. Aerial righting, landing buffering (landing with legs toward the substrate to absorb impact) and resetting (achieving a favorable body position for subsequent jumps) are crucial for a successful jump cycle [26, 27]. Moreover, eliminating the need for self-righting maneuvers after landing helps reduce predation risk, highlighting the evolutionary and functional importance of aerial righting [27].

### Wax ‘tails’ in planthopper nymphs

For planthopper nymphs, anecdotal evidence hints that they use their wax structures for gliding, yet this function remains unverified [21, 28]. Prior research on insect flight stabilization suggests that posterior filaments might assist in mid-air body pitch stabilization by generating additional drag [29]. Theoretical calculations for woolly aphids and experiments with halter-disabled fruit-flies – where attaching dandelion fibers to their abdomen improved body pitch stabilization – underscore the aerodynamic advantages wax tails could offer in planthopper nymph aerial maneuvers. As aerial dynamics play a crucial role in the survival and mobility of planthopper nymphs, unraveling the impact of wax structures on these dynamics becomes paramount.

This study sets out to assess the biomechanical role of wax tails in the post-jump aerial dynamics of planthopper nymphs. We aim to evaluate how wax influences nymph jump performance, encompassing take-off kinematics, mid-air stabilization, trajectory, and landing, by contrasting conditions with wax intact against those where it has been removed. Through in-field experimental investigations, this research explores native wax structures and their significance in insect jump performance, offering new insights into their biomechanical functions and potential evolutionary implications.

## Materials and Methods

### Study animal and environmental conditions

We collected nymphs of *Ricania sp*. from a local farm in Barangay, San Isidro, San Pablo City, Laguna, Philippines in October 2023. The site is approximately 34 km from the University of the Philippines Los Banos (UPLB) at approximate coordinates: Latitude 13.988722, Longitude 121.313120. The nymphs, found foraging on creeping cucumber, *Melothria pendula*, from the Cucurbitaceae family, were carefully collected using large plastic containers and then transported in a mesh cloth rearing cage to minimize stress. We relocated the nymphs to the University of the Philippines Los Banos - Institute of Weed Science, Entomology and Plant Pathology (UPLB-IWEP) for rearing and experimentation. In larger rearing cages, we provided the nymphs with freshly harvested stem cuttings and shoots of *M. pendula* every two days until they reached adulthood (Supplementary Video 1). The nymphal stages spanned 20-38 days before transitioning to their final morphs. After reaching adulthood, we collected, curated, and identified them as member species of the Ricaniidae family within the genus *Ricania*, using available keys [30].

All experiments took place at the farm in a semi-enclosed building or in the laboratory, both under ambient environmental conditions. The average temperature and humidity during experiments were 87.4^°^*F* ± 2.1^°^*F* and 76.3% ± 7.0%, respectively.

### High-speed videography

We recorded the rapid jumps of planthopper nymphs using a high-speed camera (Photron FASTCAM Mini AX 2000), as shown in Figure 2C, at frame rates of 2000 and 2500 fps, with exposure times set to 1/frame-rate. A Canon EF-S variable magnification 18-55mm lens was used to record full (including take-off, aerial translation, and landing) and partial jump trajectories. We conducted all experiments on the day of specimen collection or shortly thereafter, with specimens kept in rearing cages. The recordings captured jumps under two conditions: (1) with wax intact (Figure 2A) and (2) with wax removed (Figure 2B).

**Fig. 2.**
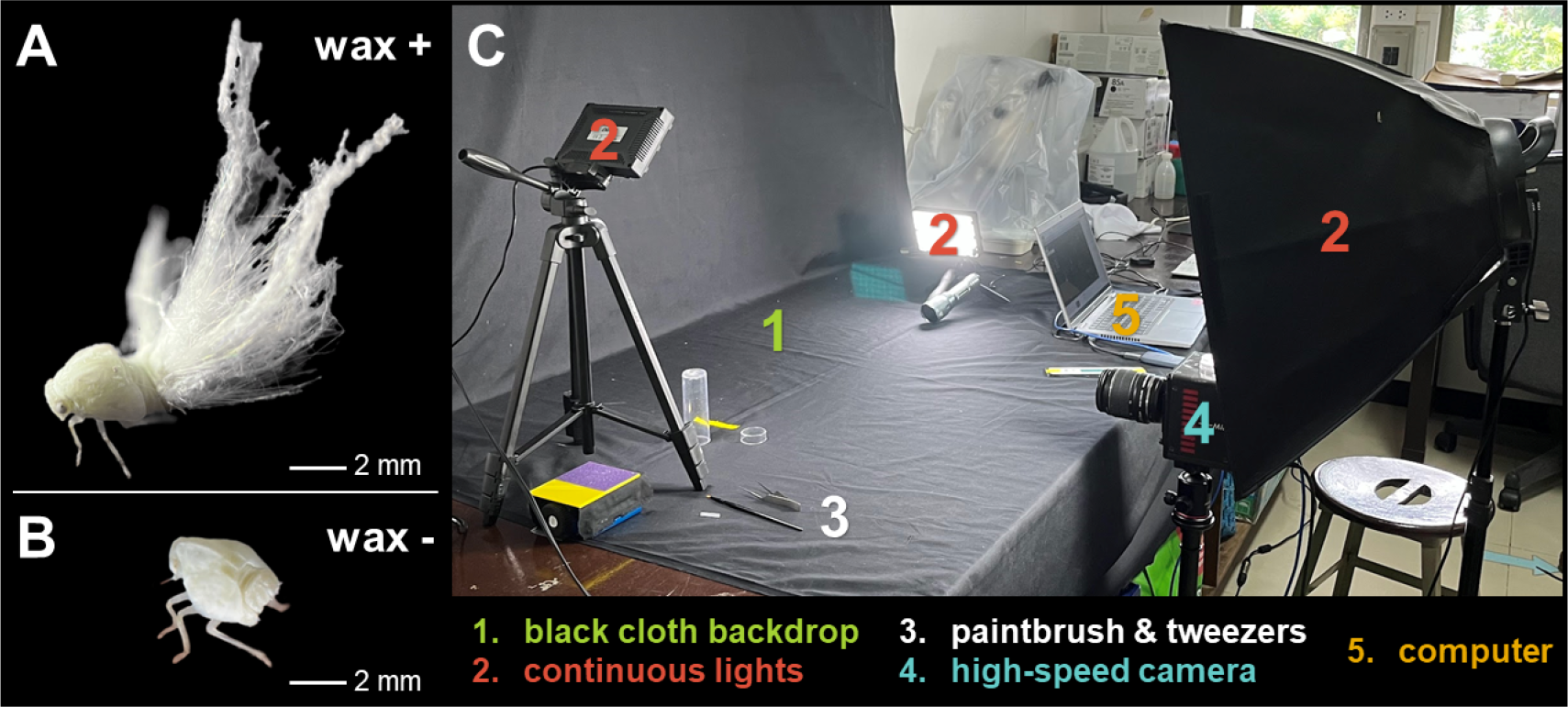
Experimental conditions and high-speed video setup: Planthopper jump conditions include (A) wax intact and (B) wax mechanically removed using a paintbrush or tweezers. (C) Experimental setup for recording jump trajectories.

The typical experimental setup included a black cloth background for contrast (Figure 2C). We recorded jumps from a cloth-covered tabletop or atop black or white paper used to guide the nymphs into frame. Continuous lighting from LED lamps mounted on tripods illuminated the area. Individual nymphs with intact wax structures were carefully obtained from rearing cages and placed in the jump arena using a 50 ml vial.

Triggering jumps involved light tapping or sliding behind the nymphs using a paintbrush, tweezers, a finger, or paper, ensuring no direct contact with the nymph or its wax. We repeated this process until we achieved desired jump recordings, as shown in Supplementary Video 2, which includes representative wax intact and wax removed jumps. Following jumps with wax intact, we used tweezers or a paintbrush to mechanically remove the wax by applying downward pressure and allowing the nymph to jump or walk away. After wax removal, nymphs rested briefly (a few minutes) before we recorded their jumps again, following the same initial procedure. Both paired experiments, where nymphs executed wax-intact and wax-removed jumps sequentially, and unpaired experiments, where nymphs jumped under only one condition, were recorded. Unpaired experiments were specifically performed to ensure fatigue did not influence wax-removed jump performance. We preserved all nymphs used in experiments in 95% ethanol for documentation and archival at the UPLB Museum of Natural History.

### Centroid tracking

We used the planthopper’s geometric centroid as a proxy for its center of mass, considering their bilateral symmetry and assuming a uniform mass density [31]. For wax removed trials, we computed the centroid from sequential binarized images from video frames, using imageJ [32]. Where image quality compromised binarization, we manually marked the centroid using the DLTdv8 digitizing software [33]. For wax intact trials, considering the wax’s mass negligible, due to its density of 0.8*g/cm*^3^, [34]) which is lower than the density of insect cuticle (1.1*g/cm*^3^, [35]), we trained iLastik, an image segmentation and analysis tool, to segment the nymph’s body from its wax filaments [36]. These segmented images were binarized using ImageJ and used to compute the centroid. We smoothed centroid positional data with a rolling 5-pt average.

We measured a maximum take-off velocity and direction during the take-off phase, designated as the moment (1 frame) just before movement onset and within 4 ms after the hindlegs lost contact with the substrate. By expressing the velocity in body lengths per second (BL/s), we ensured comparison across different individuals. This analysis covered both complete and partial jumps, including only those that fully captured the take-off phase and excluding any with slipping or significant deviation from parallel.

### Body point tracking and cumulative body rotation

We used DLTdv8 to manually or semi-automatically track two body points: (1) the perceived tip of the head and (2) the thorax-abdomen junction (Figure 3). After applying a 5pt-rolling average, we established a body axis using the tracked points and measured the body angle relative to horizontal for wax intact and wax removed conditions. We summed the changes in body angle to quantify the nymphs’ rotational behavior over time, providing a combined 2D representation of 3D rotations in pitch, yaw, and roll. This cumulative body rotation measurement represents a composite change in body angle, offering a comprehensive view of the body’s orientation during jumps. For comparison between trails, we normalized time by the total duration of the jump.

**Fig. 3.**
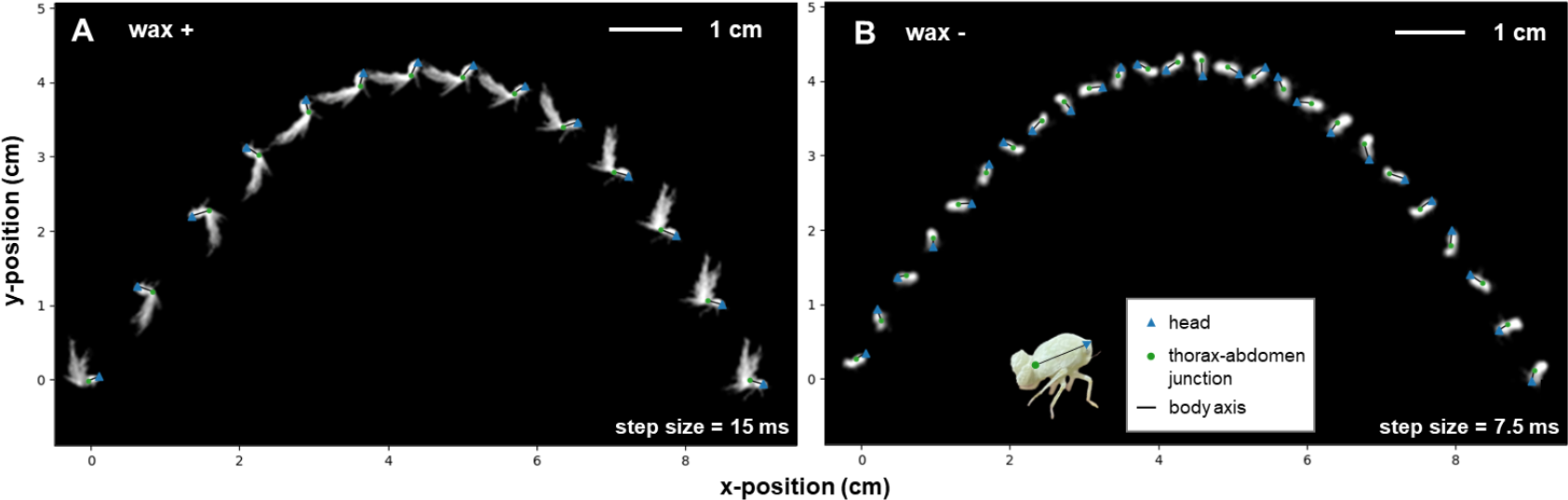
Chronophotography and digitization of planthopper nymph jump trajectories: Example planthopper jump trajectories with wax intact (A) and wax removed (B) for the same nymph. The head tip (blue triangle), thorax-abdomen junction (green circle), and body axis (black line) represent digitized and calculated positional data.

Additionally, for direct comparison of body rotations, we calculated the normalized rotation range (NRR) as shown in Equation 1.

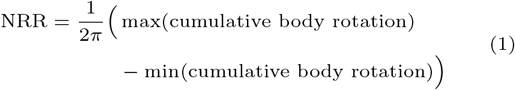

This metric, normalizing the range of cumulative body angle changes by 2*π*, scales the data to full rotations to facilitate comparison.

### Landing success

We classified a landing as successful when the planthopper touched down on its legs without rolling or bouncing after impact (Supplementary Video 3). We excluded trials where the landing was not visible or the planthopper collided with an object before landing.

### Trajectory comparison

We compared jump trajectory shapes for wax intact and wax removed conditions by normalizing centroid positions relative to the jump’s maximum length and height for x- and y-positions, respectively. We only analyzed trials that compared the entire trajectory and where the jump occurred relatively parallel to the image plane to ensure meaningful normalization.

### Statistical analysis

For jumps involving the same individual under both conditions, we used a paired t-test to assess statistical differences in take-off velocity magnitude and direction and NRR. For unpaired trials, we determined statistical difference using Mann-Whitney U tests. We applied a Chi-square test for independence to ascertain statistical significance in landing success rates.

## Results

We compared take-off velocities and directions, cumulative body rotations, normalized rotation ranges, landing success rates, and trajectory shapes between wax intact and wax removed trials to assess the impact of wax tails on planthopper nymph jump performance.

### Take-off kinematics

Paired trials revealed no significant difference in take-off velocity magnitude and direction (Table 1). Similarly, unpaired trials found no significant differences in take-off velocity, though, take-off direction showed a marginally significant difference (*p* = 0.025, **\*** *p <* 0.05). These findings suggest wax removal does not significantly affect take-off velocity, although it may slightly reduce takeoff angle. Additionally, the results from paired trials indicate that performing jumps with wax intact before those with wax removed does not significantly change take-off performance.

**Table 1.** Kinematic analysis and normalized rotation range of planthopper nymph jumps. This table presents takeoff velocity and direction, body length, and normalized rotation range for two groups: paired – nymphs executing both wax-intact and wax-removed jumps sequentially, and unpaired – nymphs jumping under only one condition. We mark statistically significant differences with **\*** *p <* 0.05, **\*\*** *p <* 0.01, **\*\*\*** *p <* 0.001. “N” denotes the number of nymphs, and “n” the total number of trials.

| Data Group Type | Wax Condition | Body Length (BL) [mm] | Take-off Velocity [m/s] | Take-off Velocity [BL/s] | Take-off Direction [degrees] | Normalized Rotation Range [rotations per jump] |
| --- | --- | --- | --- | --- | --- | --- |
| | | | Mean ± STD | Mean ± STD | Mean ± STD | Mean ± STD |
| Paired | intact (N = 1, n = 4) | 3.7 | 1.9 ± 0.3 | 511.5 ± 94.0 | 58.6 ± 11.6 | 0.8 ± 0.1 |
| | removed (N = 1, n = 4) | | 1.8 ± 0.1 | 488.1 ± 37.1 | 55.1 ± 12.6 | 6.2 ± 1.3<sup>**</sup> |
| Unpaired | intact (N = 6, n = 14) | 3.9 ± 1.0 | 2.1 ± 0.8 | 650.2 ± 266.6 | 59.2 ± 7.5 | 0.7 ± 0.2 |
| | removed (N = 6, n = 17) | 3.7 ± 0.5 | 2.0 ± 0.6 | 606.3 ± 184.0 | 51.5 ± 9.6<sup>*</sup> | 4.3 ± 1.9<sup>***</sup> |

### Aerial righting and landing success

Cumulative body rotation time series reveal striking differences in body orientation between the wax-intact and wax-removed trials. After take-off, nymphs with wax intact experience damped body angle oscillations - initial rotations followed by a counter-rotation and stabilization (Figure 4A), while nymphs with wax removed undergo continuous rotations throughout the jump (Figure 4B). On average, jumps with wax intact did not complete full rotations (NRR = 0.8 ± 0.1 rotations per jump for paired and NRR = 0.7±0.2 for the unpaired group), in contrast to jumps with wax removed, which averaged several rotations (NRR = 6.2 ± 1.3 rotations per jump for paired and NRR = 4.3 ± 1.9 for the unpaired group). The average normalized rotation range (Table 1) significantly differs between the wax-intact and wax-removed conditions for both paired (*p* = 0.004, **\*\*** *p <* 0.01) and unpaired (*p* = 1.6*e* − 5, **\*\*\*** *p <* 0.001) trials. Cumulative body rotations for unpaired jumps analyzed are shown in Supplementary Figure S1.

**Fig. 4.**
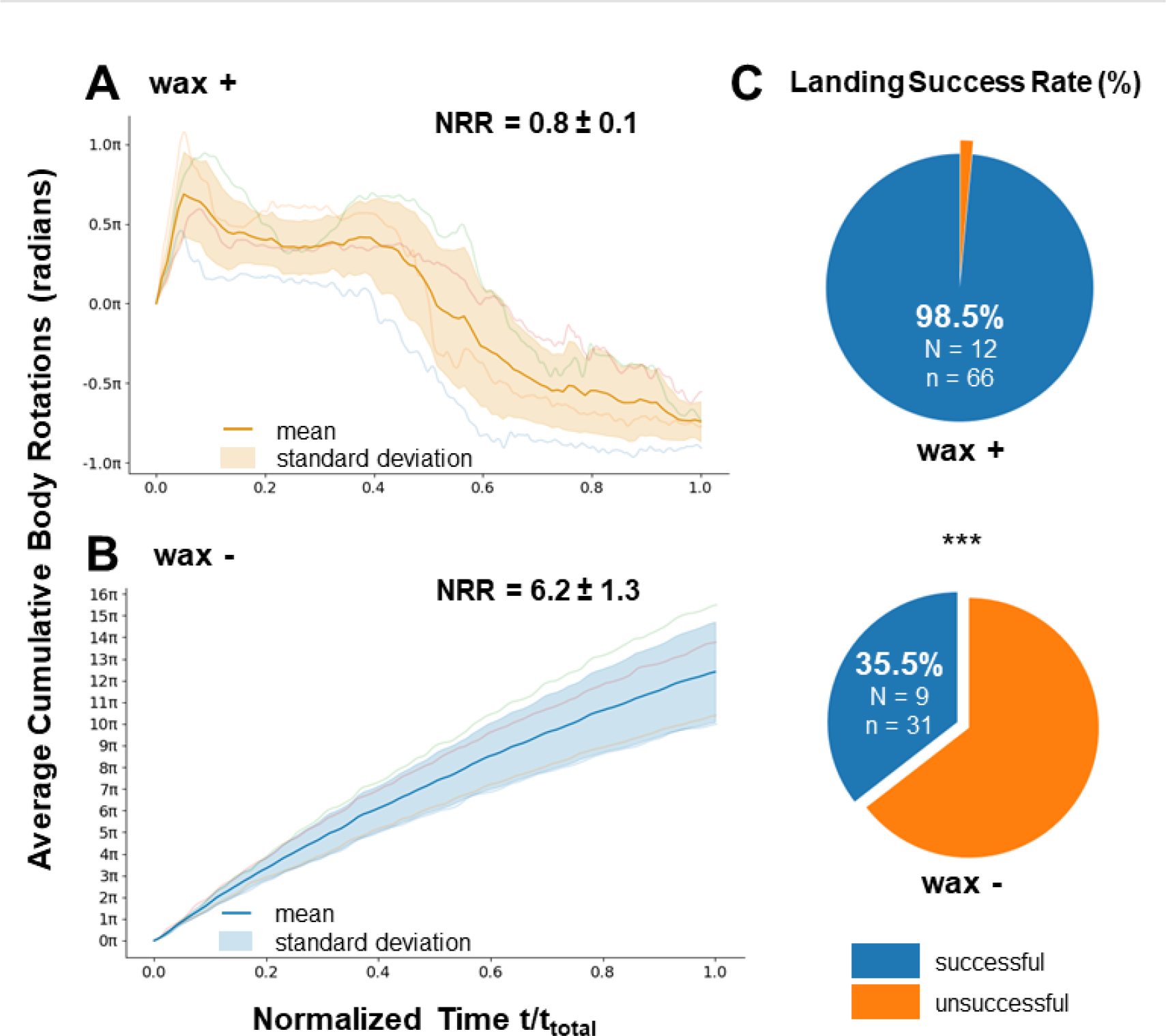
Cumulative body rotation, normalized rotation range, and landing success rate: Average cumulative body rotation and normalized rotation range (NRR) under wax intact (A) and wax removed (B) conditions for paired trials (*N* = 1, *n* = 4). Average NRR for wax intact: 0.8 ± 0.1 rotations per jump and wax removed: 6.2 ± 1.3 rotations per jump) shows a statistically significant difference (*p* = 0.004, ** *p <* 0.01); (C) The rate of successful landings for both jump conditions, with *** indicating an extremely statistically significant difference between the means (*p* = 1.4 × 10^−11^, *p* ≪ 0.001).

We note that exceptions to this pattern occurred in 2 out of the 99 trials for wax intact jumps, where nymphs underwent initial rotations post take-off. These instances had relatively low velocities (0.80 m/s and 1.35 m/s) and skewed initial take-offs (jumps towards one side, see Supplementary Video 2)

Planthoppers with wax intact successfully landed on their feet in 98.5 % (N=12, n =66) of experimental jumps, whereas wax-removed nymphs achieved successful landings only 35.5 % (N=9, n =31) of the time, significantly below a 50-50 chance (Figure 4C). This highly significant difference (*p* = 1.4*e* − 11, **\*\*\*** *p <* 0.001) underscores the critical role of wax structures in landing success for planthopper nymphs. Continuous body rotations in wax-removed jumps likely contribute to this disparity. Wax-intact nymphs often landed in a position enabling leg contact with the ground, aiding in attachment. Conversely, most wax-removed jumps resulted in nymphs hitting the ground in unfavorable orientations (i.e. contacting with the body or at an angle), leading to rolling or bouncing.

### Parabolic and Tartaglia jump trajectories

Initial conditions such as take-off velocity magnitude and direction largely determine individual jump trajectories (Figure 5, A1 and B1). However, normalization reveals the general shapes of the trajectories independent of initial take-off kinematics. Trials with wax removed follow relatively parabolic trajectories regardless of take-off velocity (Figure 5, B2). In contrast, trials with wax intact display parabolic trajectories at low velocities; however, with increased velocity, their trajectories adopt an asymmetric, triangular shape resembling a Tartaglia (Figure 5, A2) [37]. The Tartaglia trajectory, commonly observed in various ball sports such as badminton, emerges when the projectile’s velocity increases to a point where the resulting drag force exceeds the object’s weight [38, 37].

**Fig. 5.**
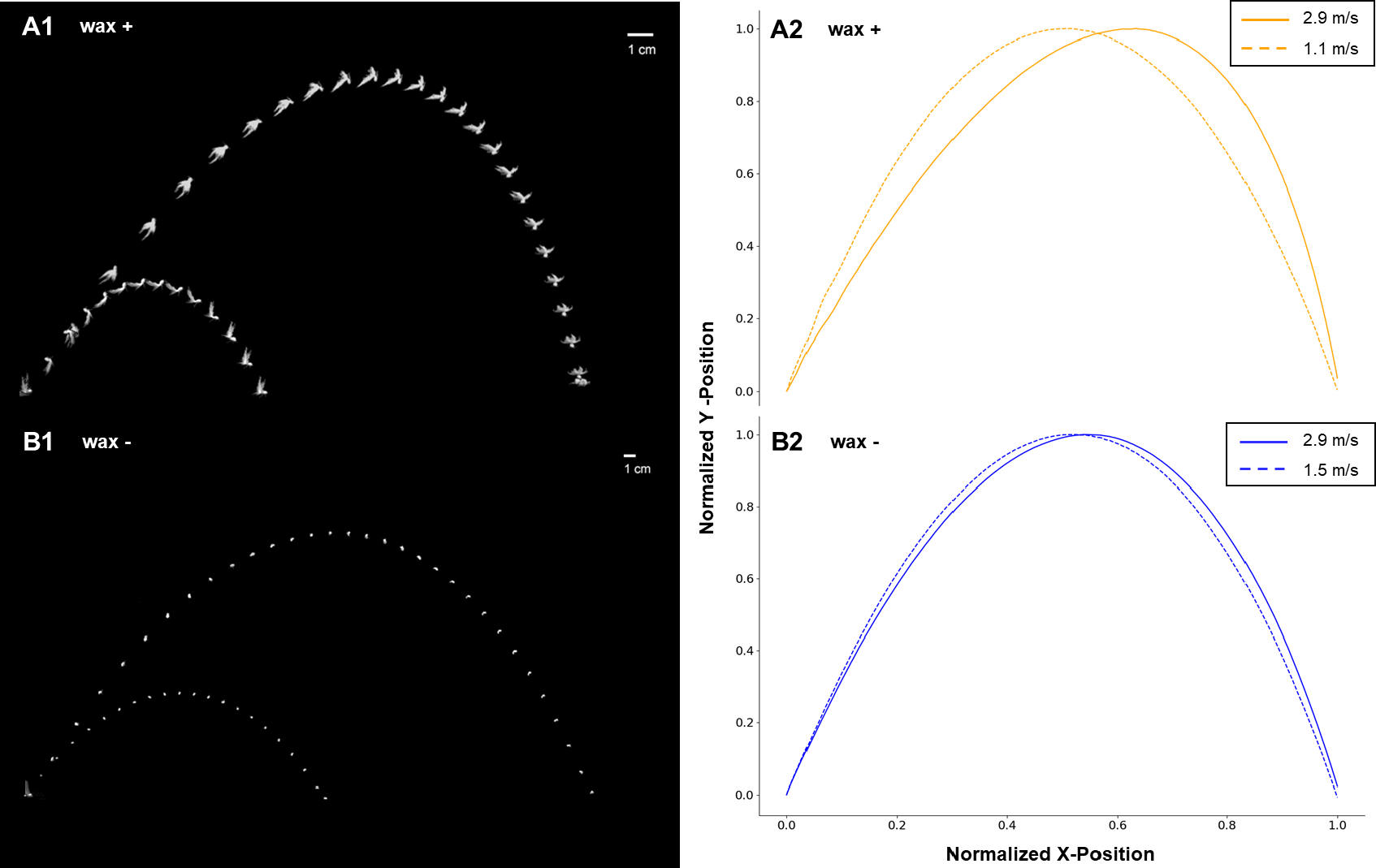
Comparison of jump trajectories with wax intact and wax removed: Chronophotographs (A1) and normalized (A2) centroid trajectories showcase jumps with wax intact at both low and high velocities. Chronophotographs (B1) and normalized (B2) centroid trajectories display jumps with wax removed at relatively low and high velocities.

## Discussion

### Jumping and aerial dynamics

Planthoppers distinguish themselves among insect taxa that produce wax structures with their exceptional jumping abilities. Over the past two decades, researchers have extensively studied planthoppers for their jump performance, power amplification mechanisms, and energy storage strategies, primarily focusing on winged adults across families such as Issidae, Dicytyopharidae, Flatidae, Derbidae, and Fulgoridae [39, 40, 41, 21, 42, 43, 44, 45, 46]. In contrast, the jumping capabilities of planthopper nymphs have received relatively less attention. Some Issid nymphs, for instance, match the jump speeds of their adult counterparts (*>* 2 m/s) [42], and specific species possess functional gears for synchronizing leg movements during jumps [47]. The persistence of jump proficiency and the evolution of specialized jumping structures in juvenile stages underscore the importance of jump performance for planthopper nymphs. Existing studies often focus on the take-off phase, neglecting critical aspects of jump performance including aerial dynamics and landing. This approach omits essential elements like aerial righting and controlled landing that are vital for executing a series of jumps in succession for escape or directed locomotion.

### Aerial righting in small jumpers

Small, wingless animals like planthopper nymphs primarily rely on aerodynamic or drag forces for aerial righting [22]. Studies on free-falling aphids, arboreal ants, arboreal bristletails, and stick insect nymphs show how appendages or passive body posture can aid mid-air reorientation via aerodynamic forces [25, 48, 49, 50]. While falling, arboreal insects may exhibit aerial translation; these instances are conditional behaviors as the insect must return to a higher altitude to repeat the action. In contrast, powerful insect jumpers, capable of repeated jumps, regularly experience significant aerial phases that necessitates stabilization strategies for successful landing. Nonetheless, research on aerial righting post-jump in small, wingless animals is still very limited. For example, semi-aquatic springtails adopt a U-shaped body posture after jumping to stabilize mid-air and land ventrally on water surfaces [51]. While planthopper nymphs and immature stages of other jumping insects, such as leafhoppers and Orthoptera (grasshoppers, crickets, and katydids), are also capable of jumping, research into their mid-air body control is lacking [52, 53]. To date, only two related studies have investigated the mid-air behavior of spotted lanternfly nymphs, uncovering and modelling their use of legs to slow rotations mid-air, which was shown to assist in landing success [54, 55].

### Wax ‘tails’ and aerodynamic forces

This study demonstrates that wax tails fundamentally alter the body orientation of airborne wax-bearing planthopper nymphs by preventing body rotations.

To understand the aerodynamic forces involved in planthopper nymph jumps, we consider the Reynolds number (Re), a dimensionless quantity that helps to compare the roles of viscous and inertial forces in flow (Equation 2).

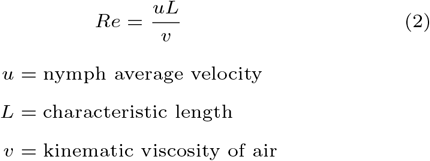

Considering wax-intact nymphs first, the characteristic length encompasses both the nymph’s body and wax tail. For these nymphs, using an average body plus wax tail length (14 mm) and velocity (2.1 m/s), we estimate *Re*_wax-on_ ∼ 2100 using kinematic viscosity for standard conditions at sea level 1.46 × 10^−5^ *m*^2^*/s* [56]. In contrast, for nymphs with wax removed, using an average body length (3.7 mm) and velocity (2.0 m/s), we estimate *Re*_wax-off_ ∼ 500. Thus, the Reynolds number for wax-intact conditions is nearly 4X higher than that for wax-removed, attributing to the extended characteristic length due to wax tails.

The finite Reynolds numbers of these animals, coupled with the intricate dynamics of their body and wax structures, complicate identifying a precise mechanism for the wax tail’s contribution to aerial righting. Unlike larger animals, such as geckos or cats, which utilize inertial forces via their tails or limbs [27], smaller organisms like dandelions, springtails, and planthoppers primarily rely on aerodynamic forces.

We hypothesize that the large surface area of the wax tails generates increased drag forces, thereby producing a stabilizing torque, similar to the effect observed in badminton shuttlecocks [38, 57] or small bioinspired jumping robots equipped with drag flaps [51]. Interestingly, it would be insightful to explore if these wax tails, owing to their porous nature, engage in unique vortex interactions similar to dandelions, where the pappus’ structure creates enhanced lift and drag forces for seed dispersal [58]. However, a comprehensive exploration of this hypothesis would necessitate particle image velocimetry (PIV) analysis, which falls beyond this paper’s scope.

The transition from parabolic to Tartaglia trajectories in planthopper nymphs reveals two distinct aerodynamic regimes influenced by their initial take-off velocity (*U*_0_) relative to their potential terminal velocity (*U*_∞_). The terminal velocity serves as a critical threshold: if *U*_0_ *< U*_∞_, the nymph experiences a classical Galilean parabola, characteristic of dense projectiles subjected to gravity and drag at high Reynolds numbers [37]. Conversely, if *U*_0_ *> U*_∞_, the trajectory shifts to a Tartaglia, characterized by a pronounced asymmetric descent that ends almost vertically, resembling an abrupt encounter with an ‘aerodynamic wall’ [59]. This phenomenon, common in sports and other projectile motions, illustrates the influence of drag forces in slowing the projectile in a manner akin to hitting an invisible barrier.

For wax-intact nymphs, the presence of wax tails potentially decreases *U*_∞_ due to additional drag, leading to Tartaglia trajectories at higher velocities where *U*_0_ surpasses *U*_∞_ (Figure 5, A1 and A2). This results in trajectories that steeply descend, facilitating controlled and stable landings – a potentially valuable adaptation for evading predators or navigating through complex environments. In contrast, wax-removed nymphs, likely having a higher *U*_∞_ due to reduced drag, predominantly follow parabolic trajectories, indicative of *U*_0_ *< U*_∞_ (Figure 5, B1 and B2). Their trajectory is often compounded by higher rotations, which further disrupt controlled descent, reflected in their significantly lower successful landing rates (Figure 4).

These findings underscore the critical aerodynamic role of wax tails in modifying the trajectory and terminal velocity of planthopper nymph jumps. While we have not directly measured *U*_∞_ for planthoppers with and without wax tails, our observations align with the expected behaviors of projectiles at high Reynolds numbers [59]. Future studies should aim to determine *U*_∞_ using wind tunnel experiments as well as validate the ecological implications of Tartaglia trajectories within the natural, unsteady flow environments where these nymphs operate.

### Limitations and future work

While the cumulative body rotation measurements help in a 2D interpretation, they do not fully capture all 3D rotations, especially out-of-plane roll rotations. Thus, it may not accurately represent the total rotational motion during jumps. Future experiments using two orthogonal high-speed cameras will better capture the 3D kinematics of the jump.

In this study, we did not assess the effect of appendages on aerial righting, although aerodynamic drag from appendages can influence body orientation, as observed in other wingless jumping insects and hexapods. We focused our analysis on the primary effects of wax removal, assuming that other stabilizing behaviors would remain unchanged across trials, as all appendages were preserved and functional in both wax intact and wax removed conditions.

Further experiments involving flow particle image velocimetry (PIV) and computational fluid dynamics (CFD) are necessary to rigorously identify the aerodynamic drag contributions from wax structures that facilitate aerial righting. Finally, implementing and validating these concepts through insect-scale robophysical models will help confirm and apply these findings.

## Concluding Remarks

Using high-speed videography, we captured the jump trajectories of planthopper nymphs with and without intact wax ‘tails’. Our findings reveal that nymphs with wax removed experience significant body rotations, whereas nymphs with wax intact exhibited rotational stability during jumps which contributed to a nearly perfect landing success rate. Calculations of Reynolds numbers and analysis of jump trajectories suggest the potential role of drag in the wax-intact system. This study is the first to demonstrate the use of wax structures for aerial righting during jump-propelled aerial translation, akin to how modifying fruit flies by attaching dandelion seed fibers enhanced their flight stability and directional control [29].

We observed similar aerial righting phenomena in nymphs of another Ricaniid species, likely *Ricania speculum*, whose wax is depicted in Figure 1G, with additional images in Supplementary Figure S2. Additionally, planthopper nymphs from the family Issidae, known for their fiber-like wax filaments, splay their fans when jumping – a behavior previously speculated to deter predators (Supplementary Video 4). These observations raise questions about how nymphs with different wax tail morphologies and behaviors might leverage aerodynamic interactions to both actively and passively control their aerial maneuvers, offering a deeper understanding of the mechanofunctions of these enigmatic wax structures.

Aerial righting strategies, jumping, and living on elevated substrates in wingless animals are considered precursors to powered flapping flight, which developed independently in insects, birds, bats, and pterosaurs [60]. How do wax structures fit into this evolutionary narrative? A leading hypothesis for flight evolution in terrestrial insects suggests that evasive jumps initiating aerial translation may have been a critical component[60]; however, winged insects appear in the fossil record without clear transitional forms [60]. Developing a broader understanding of wax structure morphologies and their biomechanical roles could enhance our comprehension of insect adaptations, and potentially inform our evolutionary understanding of insect flight.

## Funding

M.S.B. acknowledges funding support from the NIH Grant R35GM142588 and NSF Grants CAREER 1941933 and 2310691. Field work and travel for this research was supported by Georgia Tech’s Robert M. Nerem International Travel Award and Career, Research, and Innovation Development Conference (CRIDC) Travel Award. C.M. acknowledges funding support from the Georgia Tech - University Center of Exemplary Mentoring (GT-UCEM) Fellowship program and Georgia Tech’s President’s Fellowship.

## Competing Interests

No competing interest is declared.

## Author Contributions Statement

C.M conceived the research idea. G.A. R.L and C.M conducted the experiments. S.Y and G.A provided expertise on planthopper behavior, collection, and rearing. G.A identified and reared the planthoppers. S.Y. generously provided her personal farm for planthopper collection and filming. T.J and C.M analyzed the data. M.S.B and C.M interpreted the results. M.S.B. reviewed the data analysis and interpretations. All authors contributed to editing the manuscript.

## Acknowledgments

We thank members of the Bhamla and Yap labs for discussions and feedback. C.M especially thanks the De Jesus, Yap, and Laude families for their hospitality, housing, and support in the Philippines. Lastly, we thank Caroline Williams, Lisa Triedel, and Jon Harrison for organizing the symposium and the Society for Integrative and Comparative Biology for conference travel funding and the opportunity to present this research.

## Notes

### Competing Interest Statement

The authors have declared no competing interest.

